# Humans use optimal eye movements to facilitate mental rotation of objects

**DOI:** 10.64898/2026.07.02.736101

**Authors:** Emma E.M. Stewart, Ilja Wagner, Alexander C. Schütz, Roland W. Fleming

**Affiliations:** Department of Psychology, Queen Mary University London. UK; School of Biological and Behavioural Sciences, Queen Mary University London. UK; Centre for Brain and Behaviour, Queen Mary University London. UK; Justus Liebig University Giessen, Germany; General and Experimental Psychology, Phillips-University Marburg; Centre for Mind, Brain and Behaviour (CMBB), University of Marburg and Justus Liebig University Giessen

**Keywords:** eye movements, mental simulation, viewpoint perception, mental rotation

## Abstract

The ability to mentally rotate objects is a fundamental feature of human cognition, and humans can use this ability to make choices about objects based on their geometry. However, remarkably little is known about how such choices are reached, and what sort of visual information might facilitate them. We devised an experiment where participants had to mentally simulate an object’s rotation to choose which of two objects was better for a subsequent task, based on its shape alone. We also tracked their gaze while they made their choice, to see which visual information they were using to facilitate this mental simulation. We found that participants were consistently able to choose the most suitable object for the task, and, remarkably, the visual information they sampled was directly linked to their choices. Put simply, participants made better choices when they looked at more informative regions of the objects, and participants who sampled regions that were better for facilitating mental simulation made better choices overall. These findings reveal a direct link between fixations, simulation, and decision-making, suggesting that to perform any fine-grained mental simulation people need to direct their gaze at specific, informative points of an object to simulate its two-dimensional proximal image displacement.

## Main text

Humans have a remarkable ability to mentally rotate objects in their mind^1,2^, and can use this ability to make choices about objects – whether a young child deciding which shape to slot through a hole, or an adult playing “mental Tetris” to decide how to pack possessions into a small car when moving house. Such decisions require a careful mental simulation of the visual transformations each object would undergo when rotated, and then a choice of which object might be best to fulfill the current task. Despite making such seemingly trivial choices daily, it is unclear what sort of visual information is required to facilitate a dynamic mental simulation of a static physical object. Humans use eye movements to gather fine-grained visual information to make choices and decisions^3–6^, but it is unknown whether they direct their gaze to specific parts of an object to simulate its visual transformation.

We hypothesised that when faced with a task that requires simulating a static object’s rotation to make a decision – such as deciding which of two objects have a more suitable geometry for a task – participants would simulate the fine-grained visual displacement that would occur if the objects were rotated^7^. Moreover, to facilitate this mental simulation they should look at the areas of the objects that would provide the most information about this visual displacement^4,8^. We devised a task where participants were presented with the most, and least discriminable viewpoints of a single object – that is, the viewpoints that would change appearance the most or least if the object were to be rotated slightly (Figure 1A, left). They then had to choose which viewpoint they would prefer to use in a subsequent discrimination task (Figure 1A, right). Crucially this discrimination task was easier if they chose the most discriminable viewpoint rather than the least discriminable viewpoint. We predicted that to determine which viewpoint would be easier for the discrimination phase of the experiment, participants would have to simulate how much displacement would occur if each viewpoint rotated. We measured eye movements to explore whether they preferentially sampled the areas on the object that would undergo the largest visual displacement. This allowed us to test for a link between participants’ choices, fixations, and the quality of the information gathered for the mental simulation.

**Figure 1.**
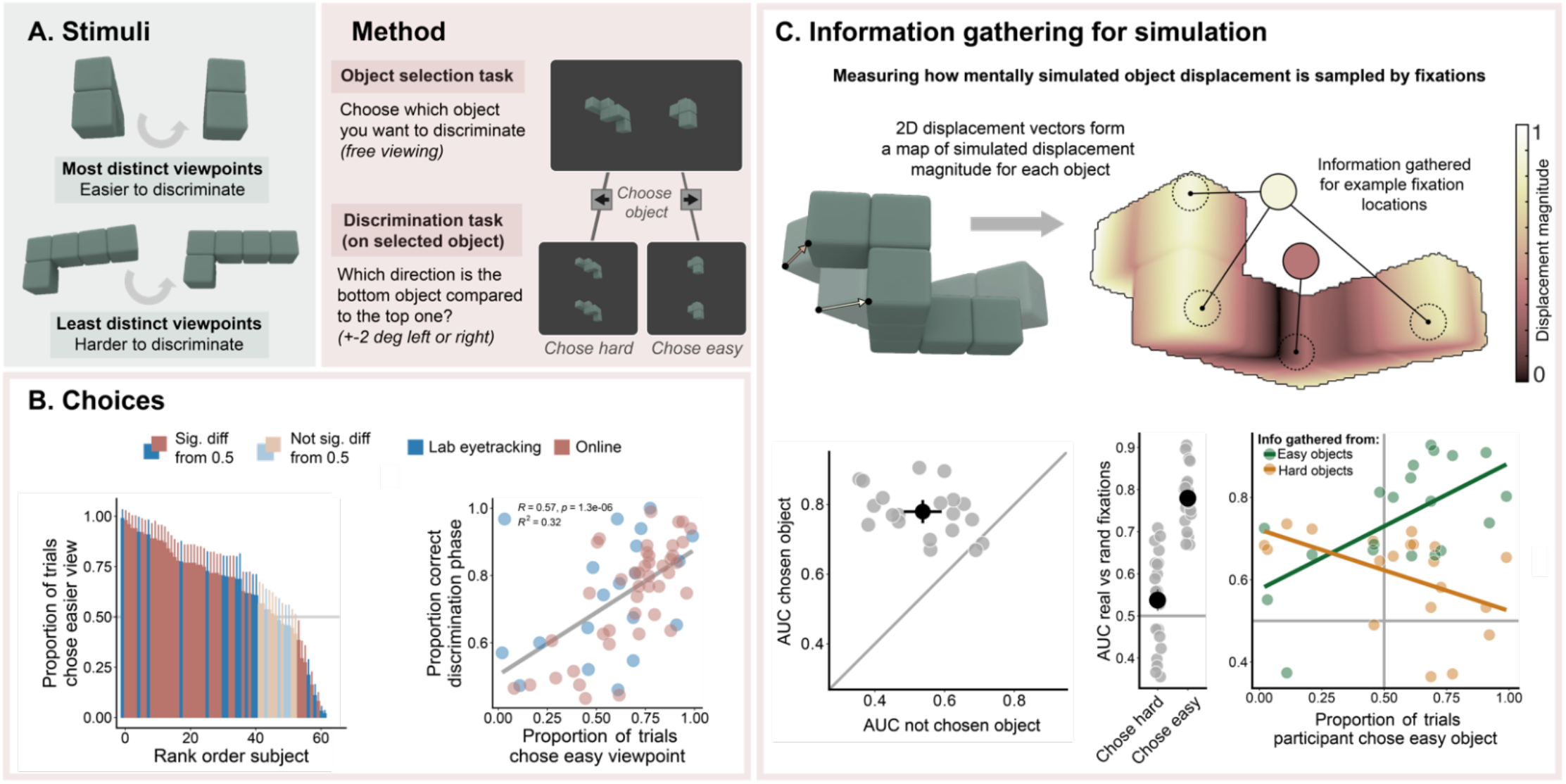
A) Stimuli and methods. Participants were shown two views of the same block-sequence object, and had to choose via button press which object would be presented for a subsequent discrimination task. In the discrimination task, the chosen viewpoint was presented in the upper half of the screen, with a viewpoint rotated ±2 degrees horizontally in the lower half. Participants were instructed to indicate via button press the direction of rotation of the bottom object relative to the top object, after which they received feedback. B) Most participants could infer the easier viewpoint (left), which was more likely to lead to correct responses in the discrimination phase of the task (right). C)Top: Using a simple optical flow model, we calculated the displacement magnitude that each viewpoint would undergo for a small rotation. For each fixation, we calculated how much displacement fell within a one-degree window of the fixation location (indicated by dotted circles). Participants sampled from more informative regions on chosen compared to non-chosen objects (bottom left), and sampled better information when they chose easy compared to hard objects (bottom middle). Bottom right: participants who were better at the choice task overall gathered better information from easy objects (green dots and line) than those who were worse at the task. For all plots, each point represents one participant.

Participants were presented with two viewpoints of a block-sequence object: the most and least discriminable viewpoints for that object, determined using a simple optical-flow model that measures the 2D displacement that would occur as an object rotates from one viewpoint to the next^7^. Participants were allowed to free-view these objects for an unlimited time, and were instructed to choose which viewpoint they wanted to use for a subsequent discrimination task. In this discrimination task, the chosen viewpoint was displayed, along with a ‘discrimination viewpoint’, which was the chosen viewpoint rotated in 3D by 2° either clockwise or anticlockwise around the vertical axis. Participants had to indicate the direction of the rotation. Crucially, this discrimination task was very difficult if the hard viewpoint was chosen, and easier if the easy viewpoint was chosen.

We conducted two versions of this experiment: 1) a lab-based study to measure participants’ eye movements during the task, and 2) an online study to assess generalisability to a larger group of participants. First we wanted to see whether participants were able to decide which was the better object and choose it for the discrimination task. Across both experiments, performance was on average better when participants performed the discrimination task on the easy compared to the hard viewpoint (Figure 1B, right, Wilcoxon Z = 1699, p <0.0001; BF_10_ = 2634, and so participants who tended to choose the easier viewpoint were more likely to be correct in the discrimination task (r = 0.57, t(61) = 5.37, p <0.0001). Participants’ decisions also reflected this performance benefit, as they were more likely to choose the easier viewpoint: 42/64 participants chose the easy viewpoint at a higher than chance level (Figure 1B) and 12 were equally likely to choose the easy or hard viewpoint.

We next wanted to investigate whether participants sampled information from regions that were the most informative about an object’s displacement to facilitate the mental simulation of object rotation. We quantified informative regions using an optical flow model, which can provide a spatial map of which regions of an object would undergo the most displacement if the object rotated. For each fixation, we calculated the maximum amount of optical flow information that fell within a 1-degree radius window around the fixation location (Figure 1C, see Supp Materials for details). We compared actual information uptake with randomly shuffled fixation and object pairs from 1000 bootstrapped samples, and used an ROC analysis to determine whether participants sampled more informative regions than would be expected by chance (AUC > 0.5). If participants require fine-grained information about specific regions of the object to perform a mental simulation in order to make a choice, we expect that participants fixate more informative regions of the object. Overall, participants sampled more informative regions than predicted by chance (AUC for information gathered across the whole trial was >0.5 for all participants). They also sampled more highly informative areas of chosen objects than they did in non-chosen objects (t(19) = -6.87, p<0.0001), suggesting a link between information gathering and choices. Participants also gathered better information in trials where they chose the easy object (Figure 1C, bottom middle; t(19) = - 6.87, p<0.0001), and individual participants who were better at the task overall (who had a higher proportion of trials where they chose the easy object) gathered better information from the easy objects than people who were not so good at the task (Figure 1C, bottom right; in a simple linear model there was a main effect of task difficulty: F(1,36) = 16.78, p<0.00023, and an interaction between object difficulty and proportion of trials in which the participant chose the easy object: F(1,36) = 16.96, p = 0.00021).

Our results suggest that people can make decisions about which object is best for a given task based on a mental simulation of the object’s 2D transformation. To make this decision, they fixate on object regions where a rotation would lead to particularly large shifts in the image. Participants whose fixation strategies target these more informative regions tend to perform better at this task. This provides evidence that humans use eye movements either to gather information in preparation for mental simulation, or while evaluating the outcomes of the simulation (or both), and that these processes directly affect their choices. This speaks to an emerging body of evidence that humans may use eye movements to simulate an object’s motion^9,10^ or predict future changes in a scene^11^, but goes beyond this to show that visual information aids the simulation of a change, such as a rotation, that a static object might undergo. It also shows a direct link between fixation, simulation, and decision-making, suggesting that to perform this fine-grained mental simulation people need to direct their gaze at specific, informative points of the object to simulate their two-dimensional proximal image displacement, rather than imagining the gist of the object’s movement as a whole^7,12^. In planning sequences of saccades to gather information for simulation, this suggests a local rather than global information maximisation approach^4,8^. Most importantly however, it shows that our mental models and internal representations and simulations are driven by the proximal information that is gathered using carefully targeted eye movements.

## Supporting information

Supplemental methods and results

## Acknowledgements

This work was funded by the European Research Council (ERC) under the European Union’s Horizon 2020 research and innovation programme (grant agreement No. 101001250 to A.C.S and Advanced Grant “STUFF”, Project No. 101098225 to R.W.F.); EU MSCA Doctoral Network “EXPLORA” (Project No. 101226908 to R.W.F.); and the Deutsche Forschungsgemeinschaft (German Research Foundation, DFG) under Germany’s Excellence Strategy (EXC 3066/1 “The Adaptive Mind”, Project No. 533717223 to A.C.S. and R.W.F) and DFG grant number 460533638 to E.E.M.S. We would like to thank Andre Gomes for help with data collection.

## Data availability statement

All experimental data, analysis materials and documentation will be made available upon publication on Zenodo (reserved DOI-http://doi.org/10.5281/zenodo.20799339).

