## Supplemental methods and results for "Humans use optimal eye movements to facilitate mental rotation of objects"

### Supplementary Material

#### Supplementary Methods

*Participants*

Eyetracking experiment: 20 participants (12 female, 8 male, mean age 26.7, SD = 4.5) completed the study for monetary reimbursement. Online experiment: after exclusions (see below), 44 participants (17 female, 1 non-binary, 26 male, mean age 29.8, SD = 8.9) were included in the study. They were recruited through Prolific and given monetary reimbursement. All participants were naïve as to the purpose of the experiment, had normal or corrected-to-normal vision, and were given monetary reimbursement. Experiments were approved by the local ethics committee of the Psychology Department at Justus-Liebig University Giessen (ethics approval number 2020-0033) and were conducted in accordance with the Declaration of Helsinki (1964).

*Equipment*

Experiments were run using custom-written software in Matlab 2020b, using the Psychophysics toolbox (Brainard, 1997; Cornelissen et al., 2002; Pelli, 1997). Stimuli were presented on 53.5x30.5cm ASUS VG248 monitor with a resolution of 1920x1080 and a refresh rate of 120Hz. Participants were seated with their heads stabilized 50cm from the monitor. Participants' eye movements were recording using an Eyelink 1000+ eyetracker, tracking monocularly at 1000Hz. Responses were collected using a standard keyboard.

*Stimuli*

Objects were generated and rendered as in Stewart et al, 2022. 100 block-sequence objects were created as 3D meshes in the ShapeToolbox[3] in Matlab. Each shape consisted between three and twelve blocks, connected in a random configuration. The objects were rendered in Blender 2.9 [4] with directional lighting coming from four directions: front (aligned with the camera), directly above the object, above and offset such that light was directed at a 45-degree angle from both the left and right. This provided full illumination of all visible sides of the cube-based objects. Objects were rendered from 360 equally-spaced viewpoints as the object rotated around the vertical axis, and were viewed slightly from below. Each object was rendered on a grey background as a 512x512-pixel image, which subtended an area of 5.5 degrees. The object occupied approximately 75% of this space. Easy and hard object viewpoints were predicted using an optical flow model that has been found to capture nonuniformities in object pose dissimilarity [1]. Easy viewpoints were the model-predicted maximally discriminable viewpoints, and hard viewpoints were the model-predicted least discriminable viewpoints.

*Procedure*

In each trial, participants were first shown the two viewpoints (model-predicted easy and hard) of a single object. The objects were presented at 3 degrees horizontal eccentricity from the screen centre, and subtended approximately 4 degrees. Participants were instructed to free-view the objects while their gaze was being recorded, and to indicate (via keypress of the left or right arrow) which object viewpoint they would like to use for a subsequent discrimination task. They were then presented with the chosen viewpoint, located 3 degrees above the screen centre. A comparison viewpoint was presented at 3 degrees below the screen centre: this view was rotated randomly ±2 degrees horizontally relative to the chosen viewpoint. Participants had to indicate (again via left or right key) whether this bottom, comparison object was rotated left or right relative to the top object. The response was untimed, and participants were allowed to free-view the objects. After responding, participants were given feedback (correct/incorrect). Participants completed 100 trials (one trial for each object), which lasted approximately half an hour.

*Procedure online study*

We conducted a follow-up online experiment which had an identical procedure to the lab-based study, with the exception that we did not record eye movements, and recorded participant responses on the Vividness of Visual Imagery Questionnaire (VVIQ)[2] (see below).

*Exclusions*

No exclusion criteria were applied in the eye-tracking study. To ensure data quality in the online version of the study, we excluded participants with a response time less than 750ms for the choice part of the experiment, or who failed an attention check question during the VVIQ phase of the experiment (6 participants excluded in total).

*Eye movement analyses*

Saccades were classified offline using the Eyelink 1000+ algorithm (velocity threshold = 22 deg/s, acceleration threshold = 3800 deg/s$^2$). Datapoints classified as blinks by the Eyelink algorithm were removed, as well as data ±30ms either side of the blink. All remaining datapoints were classified as overlapping with either the left or right object (choice) or upper or lower object (response), and were split into individual fixations by separating periods of fixation data that were separated by more than 10ms. To determine the direction of the causal relationship between fixation and choice, we divided the fixations in each trial into early (first half of total fixations) versus late (second half of total) fixations (for example in a trial with 8 fixations, the first 4 fixations would be considered early, the second four late).

*Information uptake analysis*

For each object viewpoint, a 512x512 matrix was computed which represented the optical flow information for each pixel of the object (0 where there was no object). These values were normalised between 0 and 1 for each viewpoint. Information uptake was measured as the maximum optical flow information that fell within a 1-degree radius circular window around the mean eye position for each fixation. The maximum information gathered on each trial was then recorded separately for each object. To calculate the AUCs, the maximum information per object was compared with the maximum information gathered from a shuffled control. Over 1000 samples for each participant, fixations from a random trial were paired with a randomly selected object, and the maximum information from this shuffled pair was computed. For the ROC analysis, we calculated the probability of a participant gathering more information than a set of threshold values (0 to 1 in steps of 0.01), for the real compared to the randomly sampled fixations (following the method used by [3]). The AUC was then computed for each participant.

*VVIQ questionnaire*

In the online version of the study, each participant completed the Vividness of Visual Imagery Questionnaire (VVIQ) [2]. This questionnaire has 16 items, which can be rated on 5-point scale ranging from "No image at all, you only "know" that you are thinking of an object" to "Perfectly clear and as vivid as normal vision". The scale items can be broken down into further subscales, for example Movement Imagery, which involves imagining movement, for example "You enter the shop and go to the counter. The counter assistant serves you. Money changes hands.". The questionnaire was scored such that 1 = no imagery and 5 = full imagery.

*Statistical analyses*

All statistical analyses were conducted in R. To determine whether people chose the easier viewpoint more or less than chance used a two-sided exact binomial test for each participant. T-tests used to assess differences in AUC were corrected for multiple comparisons using a Holm correction. A paired-sample Wilcoxon test was used where assumptions were violated for comparison of early versus late fixations.

### Supplementary results

*Mental imagery ability and choice*

Here we report the results of part of the online experiment where we assessed participants' mental imagery ability, and relate it to their choices. Scores on the whole VVIQ did not correlate with viewpoint choices ($R = -0.27$, $p = 0.077$), but scores on the movement subscale of the VVIQ did correlate ($R = -0.33$, $p = 0.032$). Surprisingly, this hints that participants who had better movement imagery chose the easy viewpoint less often. While this may seem counterintuitive, it suggests that participants with better movement imagery were more likely to choose the harder objects. This may be because better mental imagery abilities made the difficulty difference between viewpoints smaller for those with better mental imagery, however the effect is small ($R^2 = 0.11$), and our sample also didn't capture any participants with very low or no mental imagery. These results are inconclusive, but we report them for the sake of transparency in our full research design.
